# Simultaneous Measurement of Gross Oxygen Evolution and Underlying Photosynthetic Redox Reactions: A Case Study Using Cyanobacteria

**DOI:** 10.1101/2024.10.03.616403

**Authors:** Oded Liran

## Abstract

In phytoplankton, the intricate balance between respiration and photosynthesis is co-regulated to ensure efficient energy management and adaptation to varying environmental conditions. In cyanobacteria, both processes occur on the same membrane, sharing electron transport carriers within the same cellular compartment. By studying the interaction between photosynthesis and respiration, we can better understand how cyanobacteria balance their energetic budget for survival. In this study, we present an integrated approach that combines tracking gas exchange between cyanobacteria and their environment with analysing the redox kinetics of the underlying photosynthetic electron transport chain. This combined system allows for real-time, simultaneous acquisition of respiration and photosynthesis data. For example, it enabled us to show that the electron transport rate generated by photosystem II, translated to *in-vivo* oxygen concentration, equals the actual concentration of oxygen produced by water splitting plus the amount of oxygen respired. We further demonstrate that our system can accurately assess light respiration in wild-type strains of cyanobacteria, which amounts to 1/10 of their photosynthetic activity under optimal growth conditions. This level of accuracy was previously achievable only with specific cyanobacteria mutants. We envision applying this system in monitoring programs to elaborate on the role of photosynthetic light reactions within the broader context of primary productivity and to understand its dynamics in response to fluctuations in external environmental conditions.

## Introduction

Primary productivity is the process of assimilation and fixation of inorganic carbon into organic matter by autotrophs (mostly phototrophs) (Falkowski et al. 1998). The main engine which is responsible for the global primary production and the associated biogeochemical carbon cycle is photosynthesis (Raven and Johnston 1991). Photosynthesis occurs on specialized membranes, termed thylakoids where these membranes harbor transmembrane pigmented-protein complexes that utilize light energy to excite electrons. The excited electrons then move through a chain of electron carriers in order to generate energy and precursors needed to the carbon fixation stage (Blankenship 2021). Phototrophic microorganisms in the oceans constitute roughly 2% of the global biomass, but their photosynthesis activity is responsible for almost 40% of the global carbon sequestration and usage, and by that they serve as the foundation of aquatic bodies food web (Falkowski 1994). Cyanobacteria are responsible for 25% of oceanic carbon assimilation and are the most dominant assemblage in oceans (Flombaum et al. 2013). Several cyanobacteria strains are able to fix nitrogen and by that obtain clear advantage over other phytoplankton in ocean oligotrophic waters (Johnson et al. 2006). For this reason, and the fact that they form the base of the food web, it is important to research their energetic budget controlled by photosynthesis and respiration.

Measuring photosynthetic productivity in water samples can be performed in several techniques that are roughly divided into two groups: measuring gas exchange between the photosynthetic microorganism and their surrounding (Burlacot et al. 2020); and measuring the spectral properties of the photosynthetic pigment-protein complexes when they are reduced or oxidized (Kolber and Falkowski 1993). The two methods are used each separately to assess primary productivity – For example out of many, (Ferrón et al. 2016) use stable isotope ^18^O labeled water to determine oxygen evolution rate, and (Gorbunov and Falkowski 2021) use chlorophylll a fluorescence in order to assess quantum yield of PhotoSystem II (PSII) as a proxy to photosynthetic electron transport rate. Combining these two techniques to simultaneously study primary productivity in photosynthetic microorganisms (algae and cyanobacteria) will provide additional information that will elaborate our understanding of aquatic photosynthesis physiology and the adaptation of these organisms to their environment (currently, there is no commercial product that can provide such simultaneous measurements).

Concurrent measurement of gross oxygen evolution and underlying photosynthetic redox reactions present a significant tool for studying photosynthesis in aquatic microorganisms, especially in cyanobacteria which are unique in a sense that photosynthesis and respiratory electron transport chain share membrane and complexes (Vermaas 2001). Concurrent measurements of both gas exchange and the underlined redox kinetics of the photosynthetic complexes will open a new dimension in understanding the multifaceted nature of the photosynthetic process. It will allow to understand not only how photosynthesis and respiration are varied simultaneously but also what were the electron transport pathways which set the acquired rates.

By using membrane inlet mass spectrometry (MIMS) (Hoch and Kok 1963), it is possible to differentiate between respiration and photosynthesis in *real-time* in light periods. This is achieved by using stable isotope ^18^O_2_ and ^16^O_2_. While ^18^O_2_ is found only in very small amount in nature, its enrichment in the sample during measurement reports on respiration during light and dark (Liran et al. 2016). When water split at the PSII site, the abundant ^16^O_2_ isotope is then evolved and PSII activity can be measured (Guy et al. 1993). Here we present a system that combines MIMS and Joliot Type Spectrophotometer (JTS) technique that allows simultaneous measurement of gross oxygen evolution and photosynthetic redox reactions using the same metabolic chamber under controlled environment. We further provide a proof of concept by comparing oxygen evolution rate measured by the MIMS to the calculated oxygen evolved out of PSII and measured by the JTS.

## Materials and Procedures

### Setup of the combined system

Standard quartz cuvette with 2 clear windows (10 mm light path) of 3.5 mL with Teflon septum plug (CotzLab, UK) is used as a metabolic chamber and is placed inside the housing of a Joliot Type Spectrophotometer (JTS-150, Spectrologix, Tenessee, USA) (Figure 1). The housing is kept at a set temperature by a circulator bath (ThermoFischer, Germany). Silicone membrane (Chen Shmuel, Israel) of 1.5 mm diameter connected to a 1/16 inch stainless steel tube serves as an entry point to the MIMS tubing system. The membrane and a connected tube penetrate the septum plug to the upper 2/3 height of the cuvette. A magnetic stirrer placed at the bottom of the quartz cuvette stirs homogenously the medium inside the cuvette.

**Figure 1.**
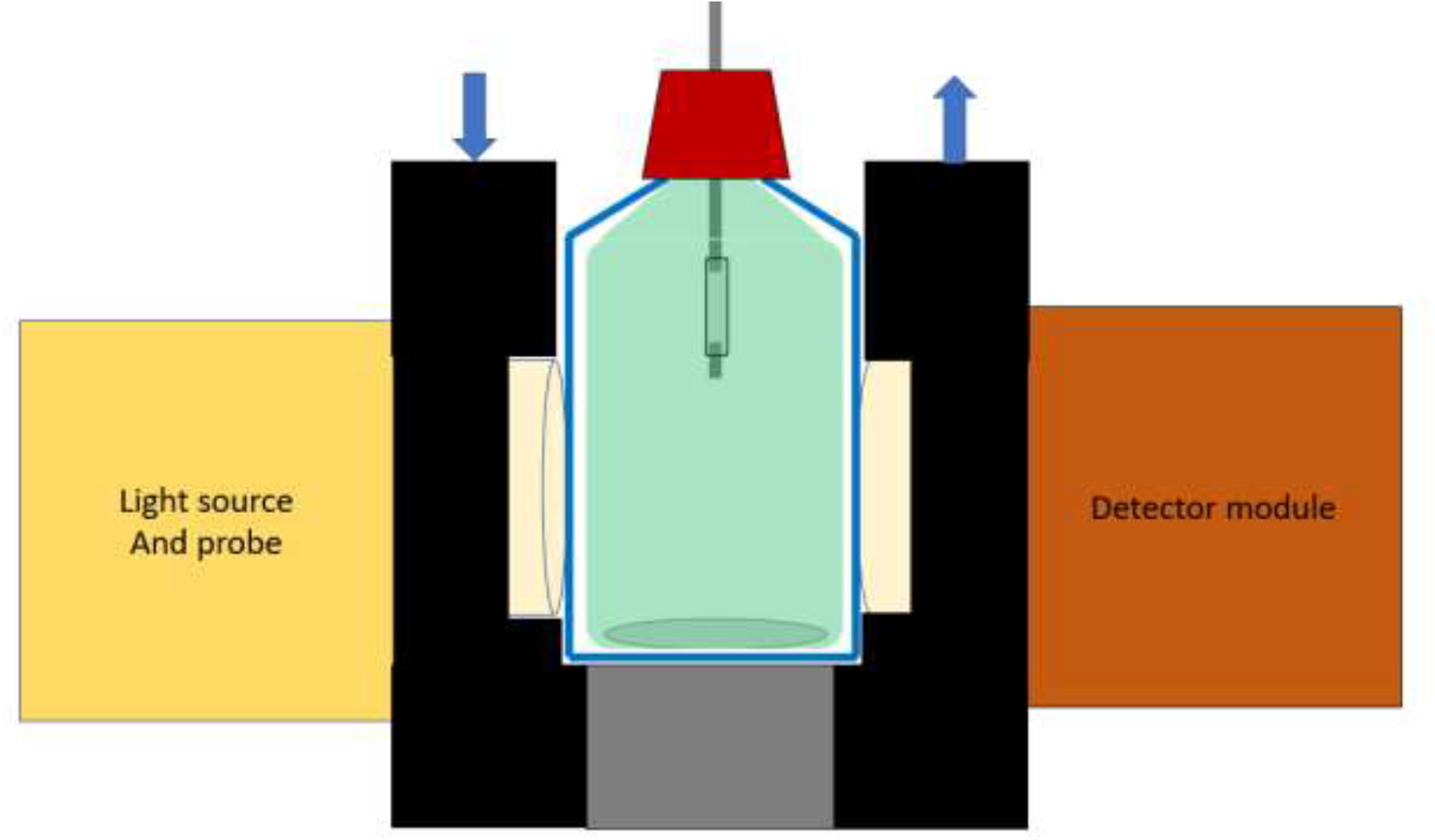
Metabolic chamber of the combined MIMS-JTS measurement system. 3.5 mL quartz cuvette (blue thin frame in the middle of the sketch) is positioned inside a chamber which is kept in a constant temperature by a water circulator (Blue arrows looking downward and upward at the left and right part of the chamber, respectively). The chamber obtains two circular opening for optics, path length 10 mm, where the light source and probe (left yellow rectangle), and detector module (right, brown rectangle) are connected to it. Magnetic stirrer is positioned within the cuvette at the bottom and controlled by a rotating magnet at the bottom of the chamber (thin grey ellipse at the lower part of the cuvette, and grey rectangle at the middle of the lower part of the chamber, respectively). Teflon screw cap is blocking gas exchange between the atmosphere and the inner part of the cuvette. MIMS nose, constructed from a silicone membrane, protrudes the septum cap into the chamber, in such a way to minimum disturb the light path of the light source.

#### Membrane Inlet Mass Spectrometry (MIMS)

MIMS (Hoch and Kok 1963) was constructed as follows: The system is divided into three parts: nose, tubing system with vacuum, and mass analyzer (Figure 2). They are connected by the tubing system. A decreasing pressure gradient is formed between these three parts in order to direct masses from the point of entry and into the analyzer. The secondary vacuum pump (Red colored square in the tubing system in Figure 2), is a rotary vane (Duo, Pffeifer vacuum, Gemerany) which lowers the pressure at the tubing system including the nose from 1000 hPa to about 2*10^−2^ hPa. The cold trap cooled down to -90^0^C in order to reduce the amount of humidity moving through the system towards the detector, and when the rotary vane is active, to minimize introduction of oil particles to the Mass Spectrometer side of the system. At this temperature, gas masses such as O_2_, CO_2_, N_2_ and Ar are free to move in the tubing system without accumulation in the trap. The primary vacuum pump (An ultra-high vacuum pump (HiCube100, Pffeifer vacuum, Germany)) reduces the vacuum pressure to ∼10^−4^ hPa on the cold trap and to ∼10^−6^-10^−7^ hPa on the entrance to the quadrupole mass spectrometer.

**Figure 2.**
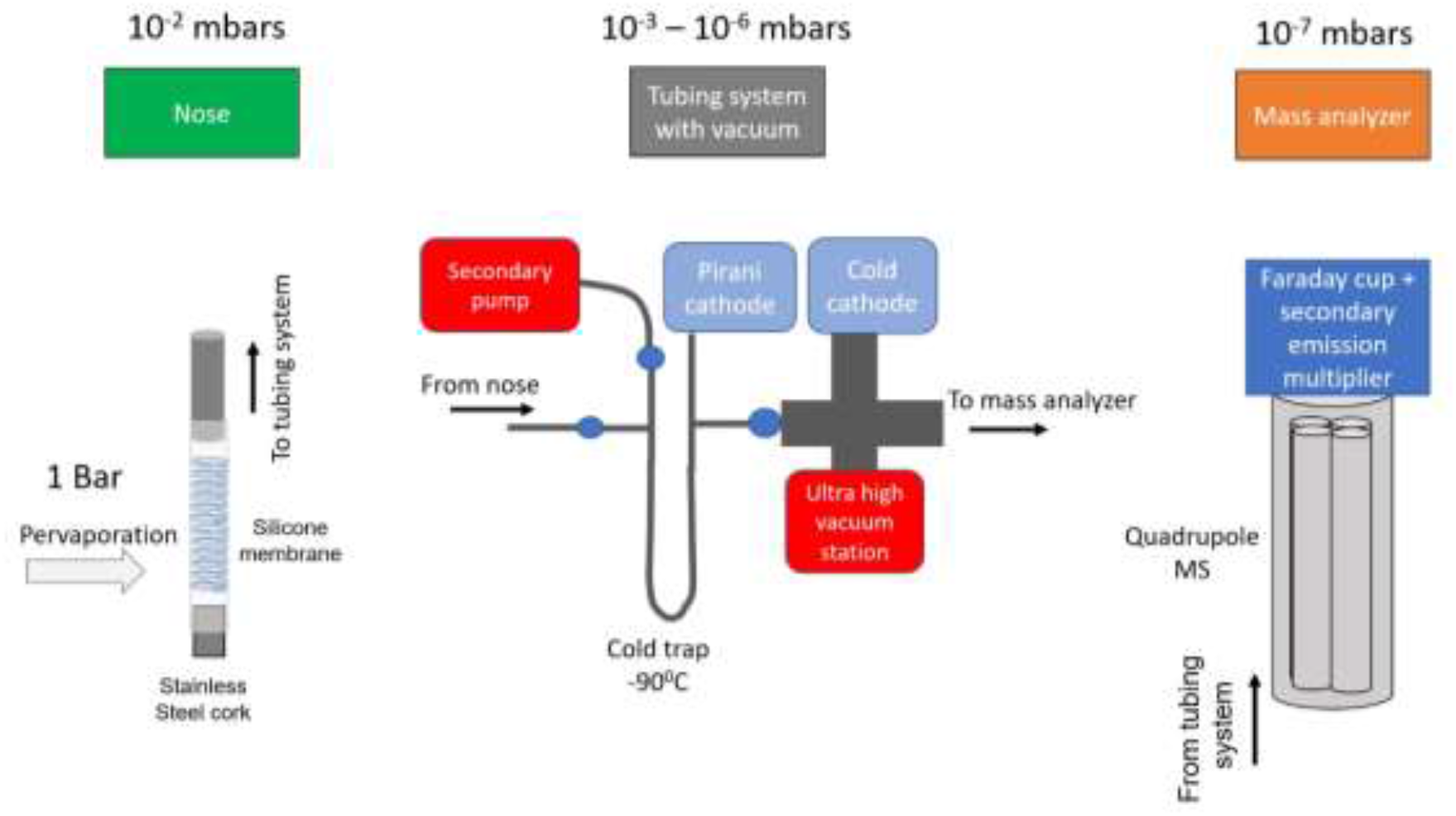
Illustration showing the three parts of a Membrane Inlet Mass Spectrometer system constructed in this study. The three parts are connected between each other with a stainless steel tubing system.

The ultra-vacuum force serves as the sink of gas masses moving from the metabolic chamber towards the detector. Therefore, the spectrometer receives only the residual gas masses that were able to escape the vacuum trap (QMS100, Pffeifer Vacuum, Germany). The quadrupole spectrometer filters only a desired gas atom with a distinct mass to charge ratio (m/z) towards a detector located at the end of the mass spectrometer. It is important to note that only at the beginning of the operation, the secondary pump is working. The primary turbopump is powerful enough to maintain the pressure gradient after it is formed within the tubing system, when one side is open to the external atmosphere at the silicon membrane nose.

#### Joliot Type Spectrophotometer (JTS)

The Joliot Type Spectrophotometer (Joliot and Joliot, 1993) is built on the assumption that very short pulses of dark period within continuous light period will not affect the kinetics of redox complexes within the photosynthetic electron transport chain (Joliot and Joliot 2002). Therefore, these periods will not affect the response of the system to external stimuli but provide a powerful glimpse to the photosynthetic machinery state during acquisition. Measuring light and saturating pulses response during these very short dark periods can report on the state of each redox complex within the Electron Transport Chain (ETC) – Photosystem II (PSII) electron transport, Photosystem I (PSI) oxidation and reduction rate, B6f complex oxidation and reduction rates, Electrochromic shift, cyclic electron transport around PSI and functional concentration of each component measured. It does so by firing a sequence of measuring light in very short light pulses at a micro second level, and increase the signal received by the use of photomultiplier units located at the measurement and reference modules (Figure 3). The reference module intercepts and records the light beam incoming into the sample, while the measurement module intercepts the light that passes the sample. In this study, the JTS was used to determine the amount of electrons generated by PSII, multiply it by the calculated PSII complexes in concentration units and divide it by four to get the amount of calculated oxygen molecules evolved from the apparatus.

**Figure 3.**
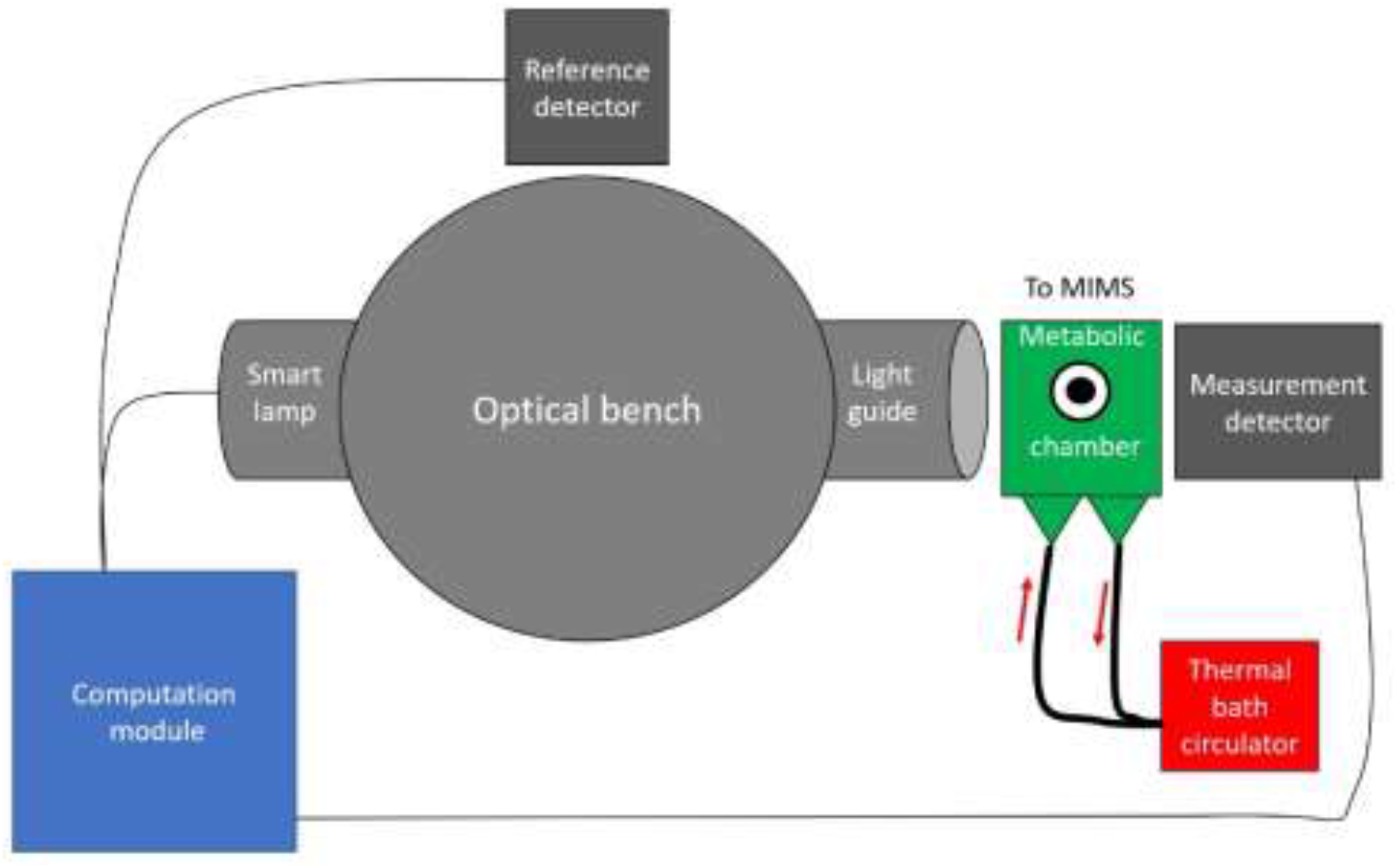
Sketch of the optical bench of the JTS system connected to the metabolic chamber.

### Materials for running experiments

BG-11 medium was prepared as suggested by (Stanier et al. 1971) with addition of 50 mM HEPES pH 7.5 (NaOH) to each sample, and 1 mM final concentration of Dissolved Inorganic Carbon (DIC) was added as NaHCO_3_ (Tchernov et al. 1997). 3-(3,4-dichlorophenyl)-1,1-dimethylurea (DCMU) and 2,5-Dibromo-6-isopropyl-3-methyl-1,4-benzoquinone (DBMIB) were added in order to block photosynthetic electron transport on PSII and B6f complexes, respectively. Stable isotope ^18^O (97%) was injected in order to calculate oxygen consumption in the light. Its concentration ranged about 1 to 5 % of total oxygen within the metabolic chamber.

### Data acquisition and interpretation in the MIMS

MIMS obtain the possibility to track the variayion in time of only desired masses, and is termed Multiple Ion Detection. In this setup the following masses were included for this study: O_2_ (Mass 32), ^18^O_2_ (Mass 36), Ar (Mass 40). Scanning of the instrument over the range of 14-44 atomic mass units (a.m.u.) was used in order to position the acquired mass over its nominal mass number. The MIMS was allowed to measure air for masses 32 and 40, and measured experimental medium with the addition of dissolved ^18^O_2_ for mass 36. The acquisition time per each mass was set to 0.5 s, with Secondary Electron Multiplier (SEM) set to 1200 V (Following setup and calibration recommended by the manufacturer). All the data was subtracted a white-noise background at a.m.u. 5.5. In order to transfer current units of the oxygen masses to concentration units, maximum solubility of oxygen was used for both oxygen masses at the temperature selected for the experiment, 20^0^C. Then, in order to subtract contribution of the MIMS equipment to the oxygen signal, Oxygen masses were normalized against Ar mass 40 as suggested by (Kana et al. 1994).

### Calculation of gross oxygen evolution and gross oxygen consumption in light and dark periods

Gross oxygen fluxes into and out of the cyanobacterial culture were calculated as previously published (Liran et al. 2016) based on the formulation of Radmer and Olinger, (1980):

a. Gross oxygen consumption in the dark: R^D^_32_
b. Net oxygen production in the light: R^L^_32_
c. Gross oxygen consumption in the light: R^L^_36_·F_32,36_
d. Gross oxygen evolution in the light: R^L^_32_ - R^L^_36_·F_32,36_

where, R is a linear regression slope extracted from the MIMS recording of oxygen concentration (µM) over time (s); upper annotation refers to the light (L) or dark (D) phase during which the linear regression slope was taken from – Dark (R^D^) or Light (R^L^); and the lowercase annotation refers to the mass measured ^16^O_2_ (R_32_), ^18^O_2_ (R_36_). F is the enrichment factor of the ^18^O_2_ isotope over the ^16^O_2_ isotope :

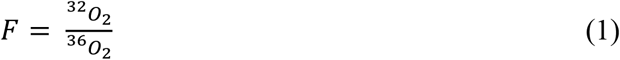

### Calculation of electron transport rates generated at PSII

PSII is the first photosynthetic complex at the beginning of the electron transport chain which is responsible to exploit absorbed light energy required to excite electrons and feed them into the electron transport chain. As PSII kinetics is measured with a fluorescence yield unit, several sub techniques based on chlorophyll a fluorescence are employed to calculate its electron transport rate in molar units. The following formulation is used:

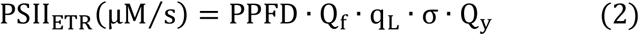

where, PPFD is Photosynthetic Photon flux Density in units of µmol photons m^-2^ s^-1^; Q_f_ is the maximum functional concentration of PSII in units of µM; q_L_ is the fraction of active PSII units out of the maximum quantity, a unitless scalar; σ is the absorption cross section of PSII in units of Å^2^/quanta of light (q); Qy is the effective quantum yield of PSII during a light period, a unitless scalar. Each acquisition of the parameters in (equation 2) is described below:

Qf [nM]: Functional concentration of PSII is calculated backwards from two parameters: the known total chlorophyll a concentration of the sample [µgr/mL] as suggested by Ritchie (2008), and the concentration of PSI [nM] calculated as described in the following sections. Essentially, each PSI monomer contains 96 chlorophylls (Jordan et al. 2001), and the molecular weight of one chlorophyll unit is 893.96 gr/mol. Therefore, the concentration of the chlorophyll could potentially be extracted from the concentration of PSI subtracted from the total calculated concentration of chlorophyll a. Then, the remaining mass has to be allocated to PSII and from that fraction PSII concentration is calculated, based on the fact that one unit of PSII contains 35 chlorophylls (Umena et al. 2011).

q_L_: The amount of active PSII complexes during a light period was calculated as suggested by Gorbunov and Falkowski (2021). The samples went through a standard induction relaxation protocol of PSII, where the sample is let dark adapted for a set period of time, and then light period follows. During the light phase the fluorescence (Ft) reaches a steady state level which is found between the Fm’, maximum fluorescence yield during light, and F_0_, the minimum fluorescence yield, just before the onset of the light period. Then, q_L_ is calculated as the following ratio:

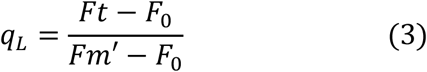

Absorption cross section (σ) (Å^2^/quantum) is calculated as suggested by (Kolber and Falkowski 1993)and fluorescence transient protocol is recorded as suggested by (Strasser et al. 2004). Then the curve is normalized at maximum fluorescence, and the region of fluorescence rise between 50 µs and 300 µs is being fitted with a double-hit Poisson function. The absorption cross section is the rate coefficient in the exponent of the Poisson function.

Effective Quantum Yield (Qy) is calculated during a standard excitation-relaxation protocol by the following formulation:

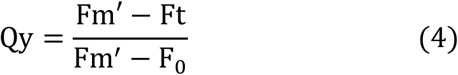

PSI functional concentration: PSI is the second photosynthetic complex capable to excite electrons by using light quanta. It is found downstream to PSII and B6f complex, where electrons moving through it will be transferred to ferredoxin to synthesize Nicotinamide Adenine Dinucleotide Phosphate (NADPH) through the Ferredoxin:NADP^+^ Reductase (FNR) complex, or will be transferred back into the electron transport chain through interaction of Ferredoxin and Photosynthesis Gradient Regulation (PGR)-like protein in complex with B6f cytochromes. In order to calculate the maximum functional concentration of PSI, two inhibitors are used – DCMU and DBMIB (Ungerer et al. 2018), and by that maximum oxidation of PSI complex is visible during a light period. Then, the maximum absorption is divided by the molar extinction coefficient of PSI which is 70 mM^-1^cm^-1^. DCMU is added in order to differentiate between the total electrons transfer through PSI and the cyclic electron transport through PSI which is not fed by PSII.

B6f complex redox rate [µM/s]: B6f complex receives electrons from the PlastoQuinone (PQ) pool through the linear electron flow, and from Ferredoxin (Fd) through a PGR-like cyclic electron flow. It donates electrons to plastocyanin (PC) and to cytochrome C6. B6f redox rate is measured by the JTS system as suggested by Ungerer et al. (2018). The redox rate is calculated as in equation 5:

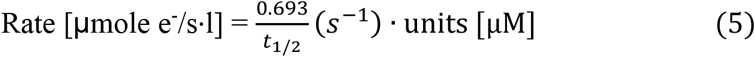

where, 0.693 is the product of ln (2) assuming a first order chemical reaction rate; t_1/2_ is the time takes the re-reduction of the complex to reach half of its maximum reduced height. Quantification of the B6f complex is performed by adding DCMU and DBMIB photosynthetic inhibitors. Maximum oxidation of the B6f complex is achieved in light, and then this value is divided by the extinction coefficient of the f cytochrome as suggested by Metzger et al. (1997).

Respiration rate before B6f complex: Succinate DeHydrogenase (SDH) receives its electrons from the Krebs cycle in the respiratory pathway and is affected by electrons filling up the PQ pool (its electron acceptors) from the photosynthetic pathway. Its activity rate *in-vivo* is assessed only indirectly by calculating the oxygen consumption by the terminal oxidase found upstream to the B6f complex which is the cytochrome BD oxidase. Therefore, oxygen consumption is measured during inhibition with DCMU and DBMIB.

## Assessment

*Microcystis aeruginosa* PCC7806 has been cultivated in BG-11 medium in 125 ml Erlenmeyer flasks (50 mL culture volume) and was illuminated by 35 µmol m^-2^s^-1^ cool day light Light Emitting Diodes (LEDs) at 20^0^C for at least five generations. Then, cultures found at mid-log phase stage were harvested for the experiment. Samples which include concentration of 10 µg (Chl a) ml^-1^ were washed and replaced with fresh BG-11 medium and were supplemented with 50 mM HEPES pH 7.5 (NaOH) and 1 mM final concentration of DIC (NaHCO_3_, Sigma, Israel). Samples were let dark adapted in the metabolic chamber for 5 minutes prior measurements. Light intensity of 1,200 µmol photons m^-2^s^-1^ was selected for the assessment step in the study by constructing preliminary light response curve (Figure 4). At light intensity of 1,200 µmol photons m^-2^s^-1^, the photosynthetic activity of PSII, starts to slow down towards a plateau, implying that some of the absorbed light energy is being dissipated by heat. Parallel measurements of oxygen evolution on the PSII side and respiration was followed.

**Figure 4.**
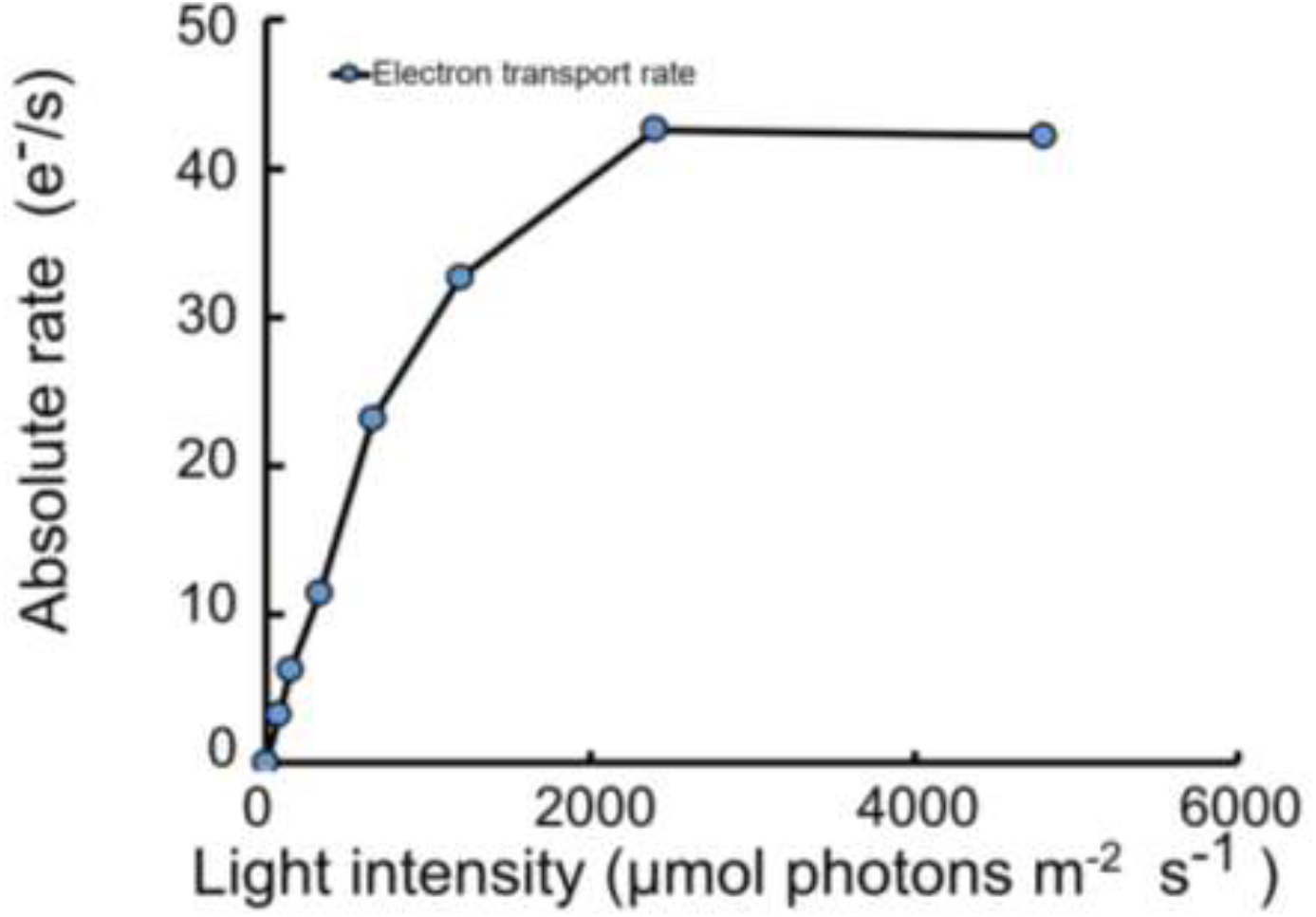
Light response curve of M. aeruginosa PCC7806 inside the metabolic chamber. Light intensities were given by the JTS module at 620 nm and absolute rate was calculated as described in the Materials and Methods.

A typical recording starts in the dark several seconds after ^18^O_2_ is injected into the sample (orange line, lower panel, Figure 5). Dark respiration is visible where ^16^O_2_ (mass 32, blue curve in Figure 5) declines in the dark. Upon switching on the light, the fluorescence trace shows sudden increase in steady state fluorescence, as expected. The gas exchange traces obtain a visible lag of about 60 s until rise in mass 32 is seen, and as documented before (Radmer and Cheniae 1976; Liran et al. 2018). Light respiration shows reaching a plateau during light and before light turned off. This is probably because ^18^O_2_ which was respired and turned to H_2_^18^O, is being split again during photosynthesis. For this reason, calculation of light respiration rate is taken from the section in the curve where ^18^O_2_ trace declines, about 100-150 s in Figure 5.

**Figure 5.**
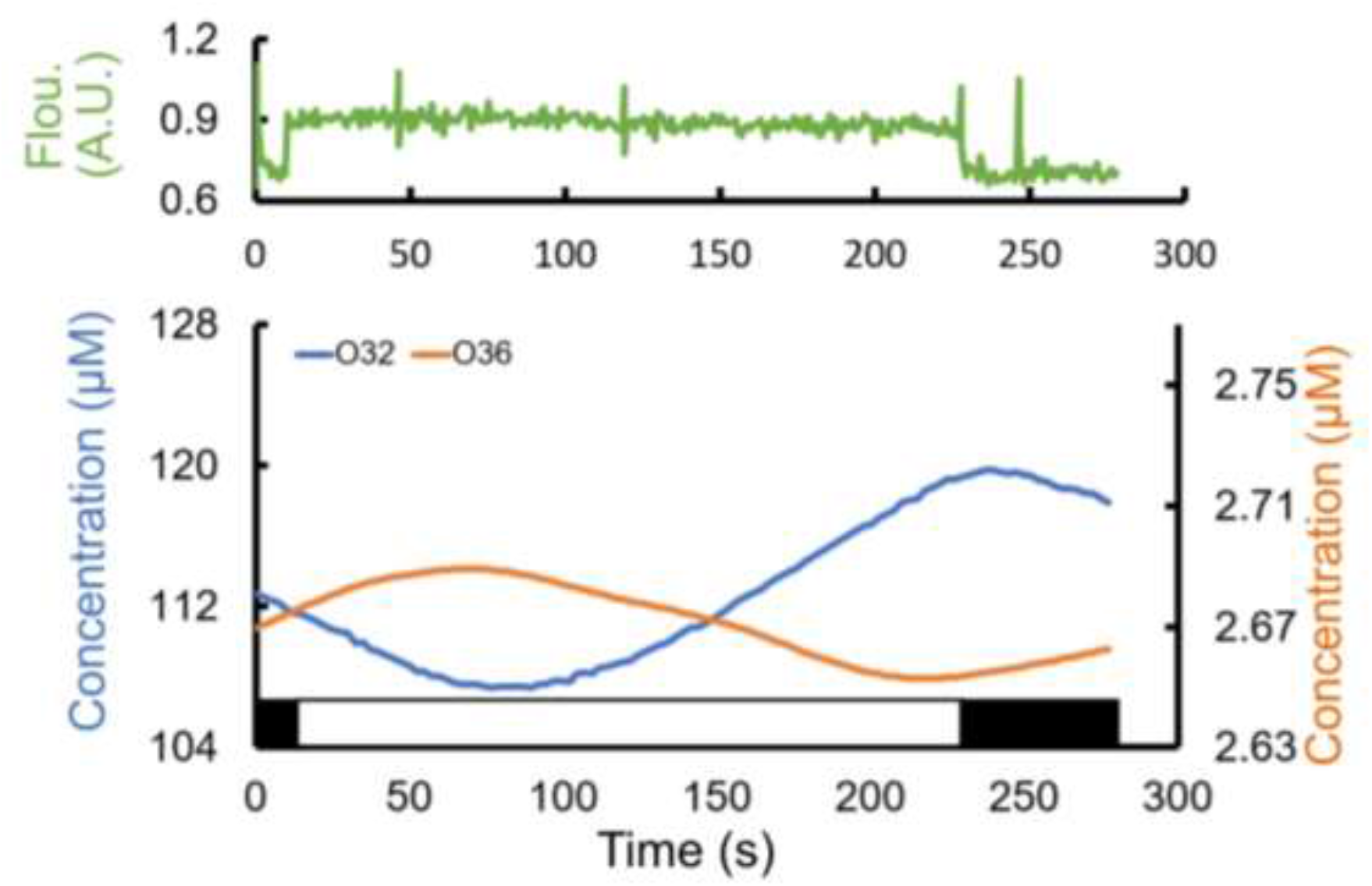
Combined Data Acquisition with MIMS and JTS. Only the relevant part of the whole gas exchange trace is shown where recording of JTS started. The figure contains two parts-The upper part describes the induction of steady state fluorescence emission (green curve) with saturating pulses along the recording; the lower part represents net oxygen evolution (blue line, O32 stands for Oxygen mass 32) and light oxygen respiration (orange line, O36 stands for Oxygen mass 36). The black and white rectangular portion of this panel represent light off and on, respectively. Figure describes one of many repeats. O36 mass line was very noisy and was smoothed using spline interpolation.

When light is switched on, PSII generates electrons that move through the the electron transport chain towards PSI. The first electron acceptor after PSII is Quinone b. Quinone b receives two electrons and becomes plastoquinone which is able to transfer the electrons on the photosynthetic route towards the B6f complex. In cyanobacteria, Quinone b is able to receive electrons also from the succinate dehydrogenase (SDH), an enzyme in the citric acid cycle (Krebs cycle). Then, it transfers it to terminal oxidase -Cytochrome BD oxidase (CYD) (Figure 6). By mesauring GOP and analyzing electron transport to either B6f or CYD we found that the electron transport rate out of PSII is 0.136 µM.By mesauring GOP and analyzing electron transport to either B6f or CYD we found that the electron transport rate out of PSII is 0.136 µM. When calculating the apporximated electron transport rate from the flourescence emission by using the JTS, we validated that the approximated oxygen eovlution rate, 0.216 µM/s is on the same magnitude and almost the same number like the actual GOP rate measured with the MIMS. When analyzing only the respiration rate while using photosynthetic inhibitor DBMIB with or without the addition of DCMU, electron transport rate downstream B6f complex is blocked. In this setup, we validated past findings about the rates of respiration and photosynthetic elelctron transport rates (Shen et al. 1993). The calculated electron transport between the SDH and the cytochrome BD oxidase was found to be about 1/10 of the photosynthetic electron transport rate as was shown in the past by using cyanobacterial mutants of PSI (Shen et al. 1993).

**Figure 6.**
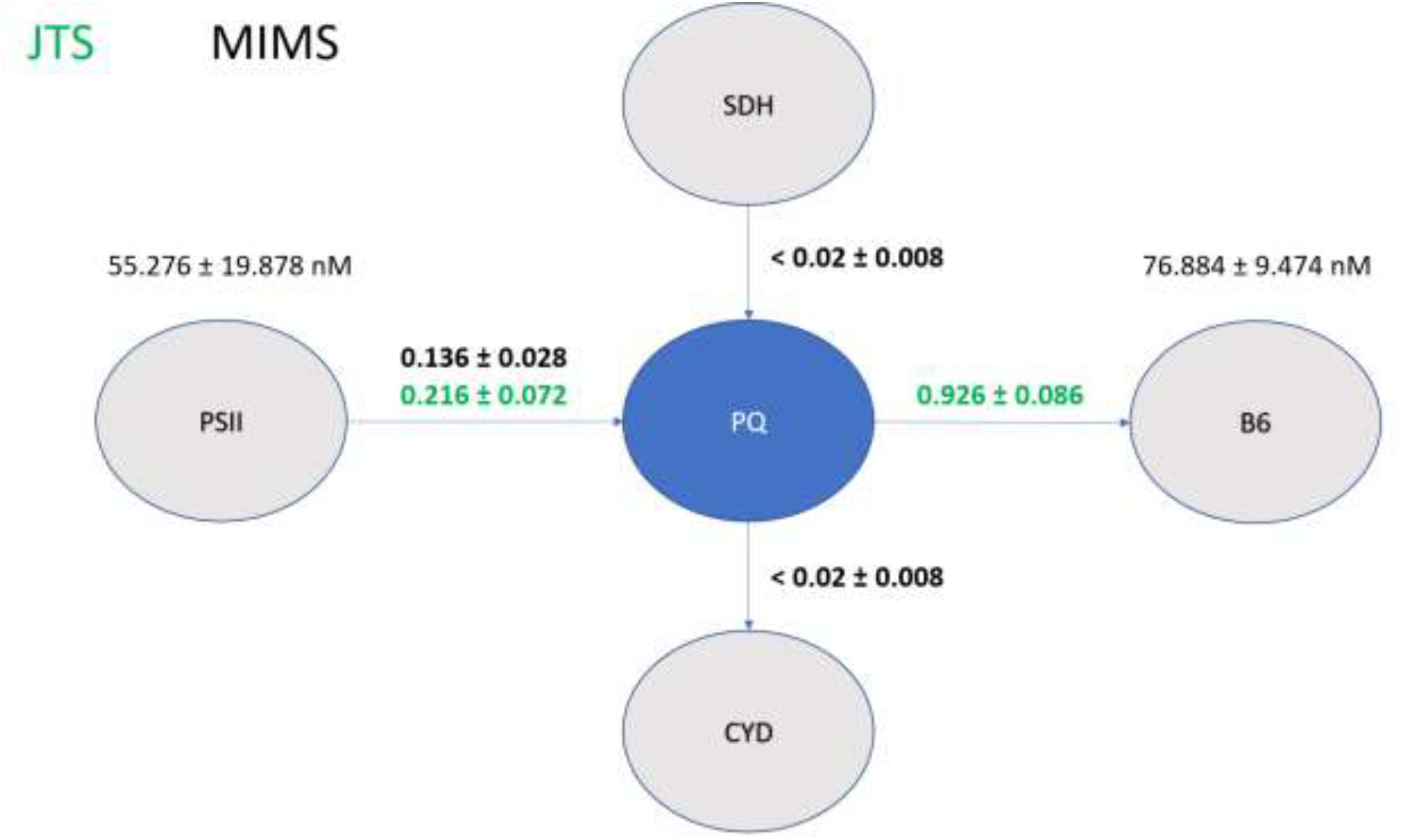
Redox rates calculation around the PQ pool junction in cyanobacterium *M. aeruginosa*. The PQ pool is shown as blue bulb in the center of the figure. The main redox complexes that donate or receive electrons from the PQ pool are shown as grey bulbs. Numbers on the arrows connecting the bulbs are electron transport rate in units of μM/s. Bold black numbers represent the actual oxygen exchanged measured and translated to electrons. The bold green numbers represent electron transport rate calculated by the JTS. Concentrations of the PSII and B6f complex are shown in numbers above the respected bulb in nM units. Error units are standard error of the mean. n=3

### Advantages and limitations in working with the combined system

The integrated system reported here obtains marked advantage over each of these systems operating separately: It is able to inspect a section within the light reactions with two different methodologies simultaneously – gas exchange (MIMS) which is verified by electron transport rate (JTS) measured by neighboring complexes, or measuring electron transport rates and verify how oxygen consumption or evolution will be interpreted. The limitations of this methodology are: a. The concentration of photosynthetic complex calculated via the maximum absorbance by using photosynthetic inhibitors refers to only the functional active units. It may not provide the actual concentration of the maximum pool of translated proteins available in the cell; b. There may be other alternative oxygen consuming complexes that may influence the rates during measurement or verification of GOP. For example, smaller participants such as the Flavoproteins which are induced only during stress (Allahverdiyeva et al. 2015) in order to reduce pressure on the two main complexes PSII and PSI, or the ARTO terminal oxidase which is present in only few cyanobacteria families (Pils and Schmetterer 2001). In such extreme experimental conditions alternative oxygen consuming complexes may obtain a significant cumulative effect that will result in large differences between the calculated electrons moving through the chain and the gross oxygen evolution calculation. One way to solve such interference will be to use a *null* mutant of either of these complexes in order to analyze its contribution to the overall respiration rate.

## Discussion

Gross Oxygen Production (GOP) is of utmost importance needed for placing primary productivity in context for operational decision making processes and development of general policies to control carbon budgets in marine environments (Sanz-Martín et al. 2019). GOP is related to carbon assimilation via the general equation of photosynthesis where one molecule of sugar will need 6 molecules of oxygen to be released versus 6 molecules of carbon dioxide to be invested. Assessing GOP in cyanobacteria poses a challenge due to the shared pool of dissolved gases in photosynthesis and respiration (Berman-Frank et al. 2003). In more details, at the compensation point, where respiration rate equals photosynthesis rate, net production is zero making it difficult to determine variations in respiration or photosynthetic activity. The proposed methodology here resolves this difficulty by measuring in parallel both consumption and generation of oxygen, thus it allows to extract the gross production even at the compensation point.

Measuring GOP as a proxy to primary productivity is considered often more accurate than the radioactive carbon tracer and fluorescence spectroscopy techniques (Sanz-Martín et al. 2019). This is because radioactive tracer technique overestimates primary production rate (Robinson et al. 2009) by assuming that all the photosynthetic energy was translated to carbon assimilation. The radio carbon tracer does not take into account respiratory steps within the light reactions before carbon assimilation (Robinson et al. 2009). In the case of fluorescence emission, not all the energy absorbed in photosynthesis is used for carbon assimilation. Part of the energy is dissipated on photoprotective mechanisms which also lower the amount of the emitted fluorescence (Muller et al. 2001). The proposed method here enables not only to record the gross oxygen evolution but also validate the calculated rate of electron transport rate from the two measurement systems (MIMS and JTS). In addition, the proposed method may report on various phenomena which is very hard to track in each method separately, for example, the cyclic electron transport around PSII (Prasil et al. 1996). Conditions of light fluctuations as may arise form turbidity developing in the water, cloudy days and light flickering in the water, may result in uncoupling of quantum yield from the gross oxygen evolution measured when measured separately on two samples. The mechanism of cyclic electron transport around PSII was found to protect algae from stress of light and by that allow them to grow faster in non-optimal environmental conditions (Ananyev et al. 2017). The proposed system may assist in determining *in-situ* occurrence of the energetic losses to primary production and by that add a correction coefficient for a better estimation of both GOP. Similar systems attempting to combine gas exchange with fluorometry, which is able to measure only net O_2_ and CO_2_ using Infra Red Gas Analyzer (IRGA) and Pulse Amplitude Modulation (PAM) were published ten years ago by Oakely et al. (2012). Their proposed system improves their model by enabling parallel measurement of both respiration and oxygen evolution due to direct discrimination of the processes by oxygen isotopes. Pradeep Patil et al. (2020) connected an oxygen optode to a cuvette provided in PAM and showed comparison between effective quantum yield and net oxygen evolution. The combined technique here improves Patil’s methodology in the fact that by incorporating a MIMS it measures gross oxygen evolution with the same units of measurement in both systems.

## Comments and Recommendations

In addition to the contribution of the combined system to basic ecophysiological research as discussed in the previous section, such a system may provide added value to monitoring programs. Current standard monitoring of primary productivity is performed using the ^14^C method (Nielsen 1952). However, this technique is dangerous and requires professional manpower to handle and perform the measurements. In addition, the ^14^C method assumes that all the ^14^C introduced to the sample is assimilated in primary production and is not used in respiration. Additionally, the technique assumes that assimilation rate of ^12^C and ^14^C are equal. These hidden assumptions result in biased measurements of primary productivity. Over the years there have been attempts to compare between the ^14^C and GOP with ^18^O stable isotope, either as dissolved oxygen or labeled water that can be split and generate labeled dissolved oxygen. Grande et al. (1989) show that the ^14^C concentration fixed equal net oxygen production without the part of respiration in the light. They concluded that during light respiration the part of ^14^C which was not fixed, was respired back to the environment during light period. Hartig et al. (1998) show that relative ETR relate to ^14^C only up until moderate light intensities, and discrepancies start at higher light intensities due to reduction in absorbed light in the algae. Laws et al. (2000) used H_2_^18^O in order to be able to measure GOP directly by labeled oxygen evolution, but found that it amounted to only half of the ^14^C assimilation rate and concluded that other consuming processes such as Mehler reaction on PSI or chlororespiration used ^18^O. Measuring *real-time* GOP with ^18^O together with pump-probe spectrophotometry may therefore provide an added value to monitoring systems. MIMS has seen great developments and adaptation when used in Ecology and Oceanography. It is used on ships to determine GOP in measurement points in the ocean (Tortell 2005), and also submersible in order to measure GOP along the water column (Bell et al. 2007). Its integration with JTS for validating GOP concentration and generation rate enhances the accuracy and reliability of primary productivity monitoring. This dual approach offers a robust validation mechanism, ensuring that the data collected is both comprehensive and precise. As such, the combined system stands to significantly advance both research and practical applications in ecological and oceanographic studies, providing a safer and a more effective alternative for monitoring programs.

## Notes

### Competing Interest Statement

The authors have declared no competing interest.

